# The histone chaperone Spt6 is required for normal recruitment of the capping enzyme Abd1 to transcribed regions

**DOI:** 10.1101/2021.05.13.444063

**Authors:** Rajaraman Gopalakrishnan, Fred Winston

## Abstract

The histone chaperone Spt6 is involved in promoting elongation of RNA polymerase II (RNAPII), maintaining chromatin structure, regulating co-transcriptional histone modifications, and controlling mRNA processing. These diverse functions of Spt6 are partly mediated through its interactions with RNAPII and other factors in the transcription elongation complex. In this study, we used mass spectrometry to characterize the differences in RNAPII interacting factors between wild-type cells and those depleted for Spt6, leading to the identification of proteins that depend on Spt6 for their interaction with RNAPII. The altered association of some of these factors could be attributed to changes in steady-state protein levels. However, Abd1, the mRNA cap methyltransferase, had decreased association with RNAPII after Spt6 depletion despite unchanged Abd1 protein levels, showing a requirement for Spt6 in mediating the Abd1-RNAPII interaction. Genome-wide studies showed that Spt6 is required for maintaining the level of Abd1 over transcribed regions, as well as the level of Spt5, another protein known to recruit Abd1 to chromatin. Abd1 levels were particularly decreased at the 5’ ends of genes after Spt6 depletion, suggesting a greater need for Spt6 in Abd1 recruitment over these regions. Together, our results show that Spt6 is important in regulating the composition of the transcription elongation complex and reveal a previously unknown function for Spt6 in the recruitment of Abd1.

## Introduction

During transcription elongation, RNAPII interacts with a large set of proteins. These include proteins that promote processivity of RNAPII such as TFIIS and the DSIF complex (composed of Spt4/5), histone chaperones such as Spt6 and FACT, histone modification enzymes such as Set2, and enzymes involved in co-transcriptional processes such as mRNA capping and splicing (1,2). The concerted action of these factors ensures efficient transcription by RNAPII and mRNA processing, producing a mature mRNA molecule that can be exported to the cytoplasm for translation. For many of these transcription elongation factors, our understanding of how they regulate transcription is limited.

The histone chaperone Spt6 is a highly conserved, multifunctional protein that is essential for viability in *Saccharomyces cerevisiae* (3,4) and in metazoans (5-7). Spt6 directly interacts with RNAPII through its C-terminal tandem SH2 domains and it is part of the transcription elongation complex (8-14). The localization of Spt6 over transcribed regions positively correlates with the level of RNAPII, suggesting an important role for Spt6 in transcription elongation, (15,16) and there is evidence that Spt6 is required for normal elongation in vivo and in vitro (17,18). Spt6 also interacts with all four histones (19,20) and it is required for chromatin organization over transcribed regions (21-24). Strikingly, in *spt6* mutants there is a genome-wide increase in the expression of transcripts originating from within gene bodies in both sense and antisense orientations (21,22,25-27). Together, these results have established that Spt6 is required for maintaining chromatin structure and transcriptional fidelity.

Spt6 regulates transcription elongation and co-transcriptional processes partly through its interactions with other proteins in the transcription elongation complex. Spt6 physically interacts with Spt5 and both proteins are required for normal transcription elongation (14,28-30). Spt6 also recruits the PAF complex (31) to transcribed regions (17,23,32), and can stimulate the ability of this complex to promote transcription elongation (14). The N-terminal region of Spt6 interacts with Spn1/Iws1 (33,34). The Spt6-Spn1 interaction is necessary for maintaining H3K36 methylation, which is co-transcriptionally deposited in yeast and human cells (35-37). In yeast, Spt6 is required for promoting the activity of the H3K36 methyltransferase Set2 (38-40). In humans, the Spt6-Spn1 interaction also promotes mRNA splicing and export (41). Given the wide range of functions of Spt6, it presents an ideal system to understand the link between transcription elongation and the regulation of co-transcriptional processes.

In this study, we set out to characterize the role of Spt6 in the transcription elongation complex using *spt6-1004*, a widely-used temperature-sensitive allele of *SPT6* (22,25). After a shift to the non-permissive temperature, *spt6-1004* cells become depleted for Spt6 protein (22) and show genome-wide reduction of nucleosome positioning and occupancy (15,21,22), loss of H3K36 methylation (38,39), and expression of intragenic transcripts (21,22,25-27). To comprehensively characterize the set of proteins whose interaction with RNAPII is Spt6-dependent, we compared the RNAPII-interacting proteins between wild-type and *spt6-1004* cells. We identified 58 proteins that have altered association with RNAPII in *spt6-1004*, including decreased association of Spt5, Spn1, and PAF complex subunits, indicating a central role for Spt6 in regulating the composition of the transcription elongation complex. Interestingly, we find that the interaction of the mRNA cap methyltransferase Abd1 with RNAPII is decreased in *spt6-1004*. However, the other two capping enzymes, Cet1 and Ceg1, have unaltered interaction with RNAPII, suggesting a role for Spt6 in specifically mediating the interaction of Abd1 with the RNAPII elongation complex. ChIP-seq analysis shows a genome-wide decrease in Abd1, Spt5, and RNAPII levels on chromatin in *spt6-1004*. The decrease in Abd1 occupancy is particularly apparent at the 5’ ends of genes, which might be due to reduced Spt5 binding at the same region. In summary, our results provide new insights into the role of Spt6 in mediating interactions of other factors with RNAPII during transcription elongation.

## Results

### Purification of RNAPII complexes from wild-type and *spt6-1004* cells

To identify factors whose interaction with RNAPII is dependent on Spt6, we purified RNAPII complexes from wild-type and *spt6-1004* cells using BioTAP-XL – a two-step purification method originally developed for *Drosophila* cells (42) and adapted here for *S. cerevisiae* (Figure 1A). To do this, we fused the C-terminal end of Rpb3 (a subunit of RNAPII) to a tandem affinity tag consisting of Protein A and a protein sequence that can be efficiently biotinylated *in vivo*. Wild-type and *spt6-1004* cells expressing the tagged *RPB3* gene were grown at 30°C and shifted to the non-permissive temperature (37°C) for 80 minutes prior to formaldehyde cross-linking and cell harvesting. This temperature shift leads to depletion of Spt6 protein in *spt6-1004* cells (Figure 1B) and exacerbation of mutant phenotypes without impairing the viability of the cells (22). Following cross-linking of protein complexes and cell lysis, a two-step purification of Rpb3 was performed, first using IgG, which binds to protein A, and second using streptavidin which binds to biotin (Figure 1A). A representative silver stained gel showed that the purification of Rpb3 from *spt6-1004* cells was dependent upon the tag (Figure 1C). As a positive control for the purification of RNAPII associated factors, we tested for the co-purification of Spt5 with Rpb3.

**Figure 1.**
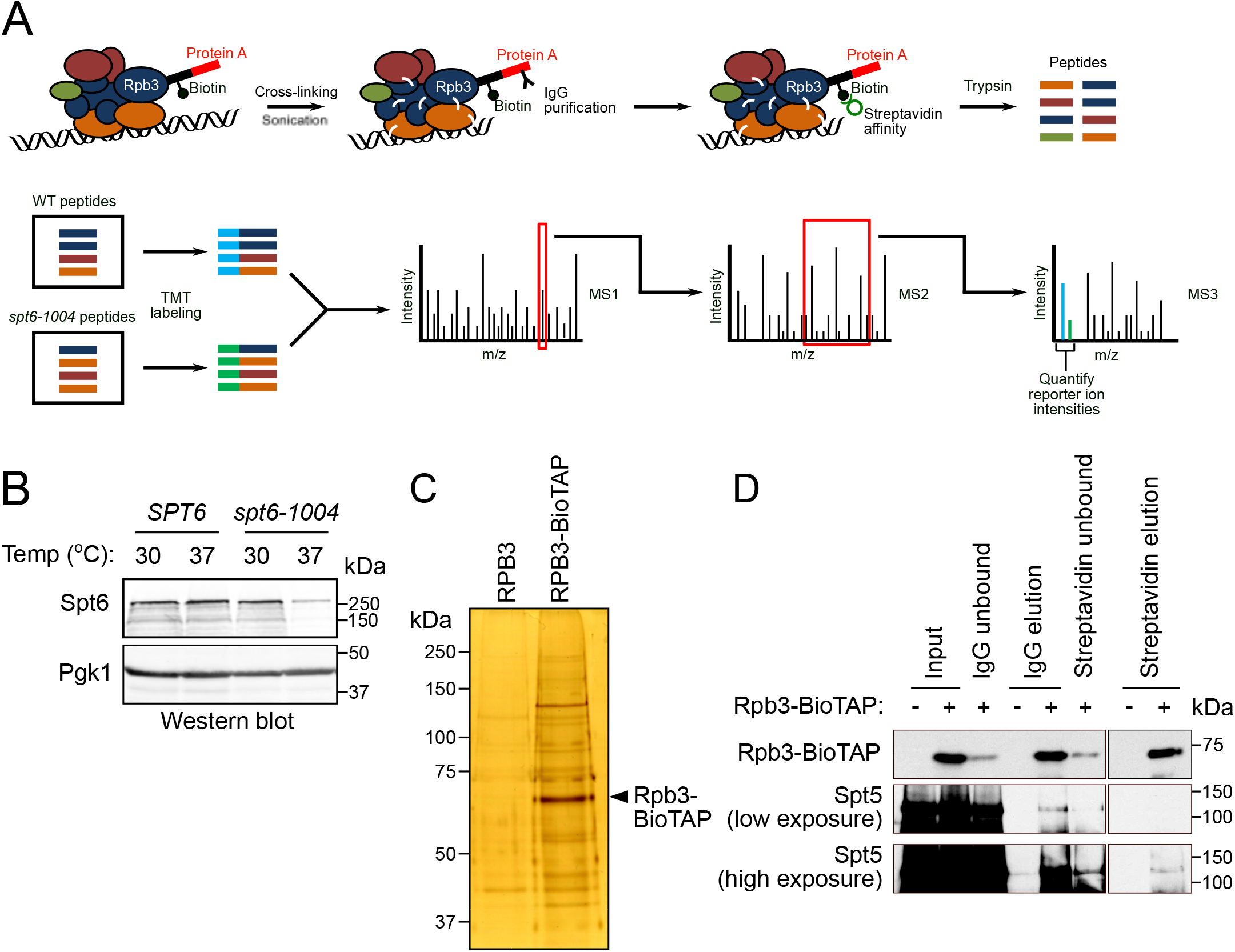
Purification of RNAPII complexes by BIOTAP-XL. (A) Schematic showing the BioTAP-XL procedure followed by mass spectrometry. The cross-linked protein complexes are sonicated to solubilize chromatin-bound proteins. The protein complexes are purified using a two-step process - first using IgG that binds to the protein A portion of the tag on Rpb3, and then using streptavidin that binds to the biotinylated portion of the tag. The purified complexes are subjected to on-bead trypsin digestion. The eluted peptides from the different samples are labeled with isobaric tandem mass tags and analyzed by triple-stage mass spectrometry. The peptide abundances are obtained by quantifying the reporter ion intensities for each sample at the MS3 stage. (B) Western blot showing Spt6 protein levels in wild-type and *spt6-1004* cells before (30°C) after after a temperature shift to 37°C. Pgk1 served as a loading control. (C) Representative silver stained gel showing purification of Rpb3 from *spt6-1004* cells with and without the BioTAP tag on Rpb3. (D) Western blot showing the abundance of Rpb3 and Spt5 at the different stages of purification.

Western blots showed that Spt5 co-purified with Rpb3 only in purifications done from Rpb3-tagged cells (Figure 1D).

### Identification and comparison of RNAPII interacting proteins in wild-type and *spt6-1004* cells

To identify the proteins associated with RNAPII in wild-type and *spt6-1004* cells, we analyzed purified Rpb3 complexes by mass spectrometry. As a control for specificity for association with RNAPII, we also analyzed proteins purified from untagged wild-type and *spt6-1004* cells. All samples were prepared and analyzed in duplicate. Peptides from each sample were labeled with tandem mass tags (TMT) to permit multiplexing and quantitative comparisons of protein levels between samples (43). The reporter ion intensities for each peptide were quantified at the MS3 stage (Figure 1A), which provides greater accuracy due to lower contamination from other peptides having a similar m/z ratio as compared to quantification at the MS2 stage (44). Averaged peptide intensities for each protein correlated well between the two replicates for each sample (Supplementary Figure 1).

To identify Rpb3-associated proteins that were specifically depleted or enriched in *spt6-1004* cells, the mass spec results were compared using Perseus (45). First, protein abundances from Rpb3-tagged wild-type and Rpb3-tagged *spt6-1004* cells were each compared to the corresponding abundances from the untagged cells. Using a permutation based false discovery rate (FDR) cutoff of 0.05, we identified 473 and 401 proteins (with more than one peptide detected for 352 and 308 proteins) that were enriched in Rpb3-tagged wild-type and Rpb3-tagged *spt6-1004* cells, respectively (Figure 2A). Second, we compared protein abundances from Rpb3-tagged wild-type and Rpb3-tagged *spt6-1004* cells to identify proteins that were specifically enriched or depleted in *spt6-1004*, and identified 137 such proteins (FDR < 0.05). Of these, 58 proteins were also enriched over Rpb3-untagged samples based on our initial comparisons (Figure 1C, Table 1). The remaining 79 proteins likely represent abundant non-specific interactors, whose gene expression may be altered in *spt6-1004*.

**Table 1.** List of differential interactors identified by BioTAP-XL.

| Proteins enriched in wild-type |  | Proteins enriched in *spt6-1004* |  |
| --- | --- | --- | --- |
| Protein | Fold change<br>(WT/*spt6-1004*) | Protein | Fold change<br>(WT/*spt6-1004*) |
| Transcription elongation factors |  | mRNA 3' end processing and export |  |
| Spt6 | 9.99 | Gbp2 | 0.43 |
| Spt5 | 1.69 | Yra1 | 0.54 |
| Spn1 | 7.45 | Yra2 | 0.49 |
| PAF complex subunits |  | Sub2 | 0.65 |
| Ctr9 | 2.58 | Hrp1 | 0.55 |
| Rtf1 | 2.47 | rRNA processing |  |
| Paf1 | 2.89 | Nog1 | 0.57 |
| Cdc73 | 2.92 | Nug1 | 0.54 |
| Leo1 | 2.87 | Erb1 | 0.63 |
| General transcription factors |  | Nop6 | 0.66 |
| Tfb1 | 1.82 | Nop1 | 0.6 |
| Tfb4 | 1.73 | Rrp5 | 0.54 |
| Tfb2 | 1.77 | Dbp2 | 0.46 |
| Toa1 | 1.7 | Nsr1 | 0.59 |
| Toa2 | 1.61 | Tsr1 | 0.38 |
| Mediator complex subunits |  | Ebp2 | 0.59 |
| Srb7 | 2 | Rpl8B | 0.5 |
| Med8 | 2.09 | Others |  |
| Cse2 | 2 | Bur2 | 0.65 |
| Med11 | 1.75 | Chd1 | 0.6 |
| Srb6 | 2.01 | Tra1 | 0.53 |
| Others |  | Nqm1 | 0.19 |
| Abd1 | 1.8 | Rep1 | 0.34 |
| Wtm1 | 2.97 | Rep2 | 0.36 |
| Emg1 | 1.63 | Orc4 | 0.45 |
| Set2 | 1.84 | Sis1 | 0.65 |
| His5 | 1.67 | Puf6 | 0.64 |
| Tal1 | 2.65 | Rex3 | 0.59 |
| Ubp14 | 1.99 | Tfc7 | 0.53 |
| Iwr1 | 3.04 | Gga1 | 0.4 |
| Utp25 | 2.24 | Trm2 | 0.42 |
| Hpm1 | 2.87 |  |  |
| Erg27 | 2.71 |  |  |

**Figure 2.**
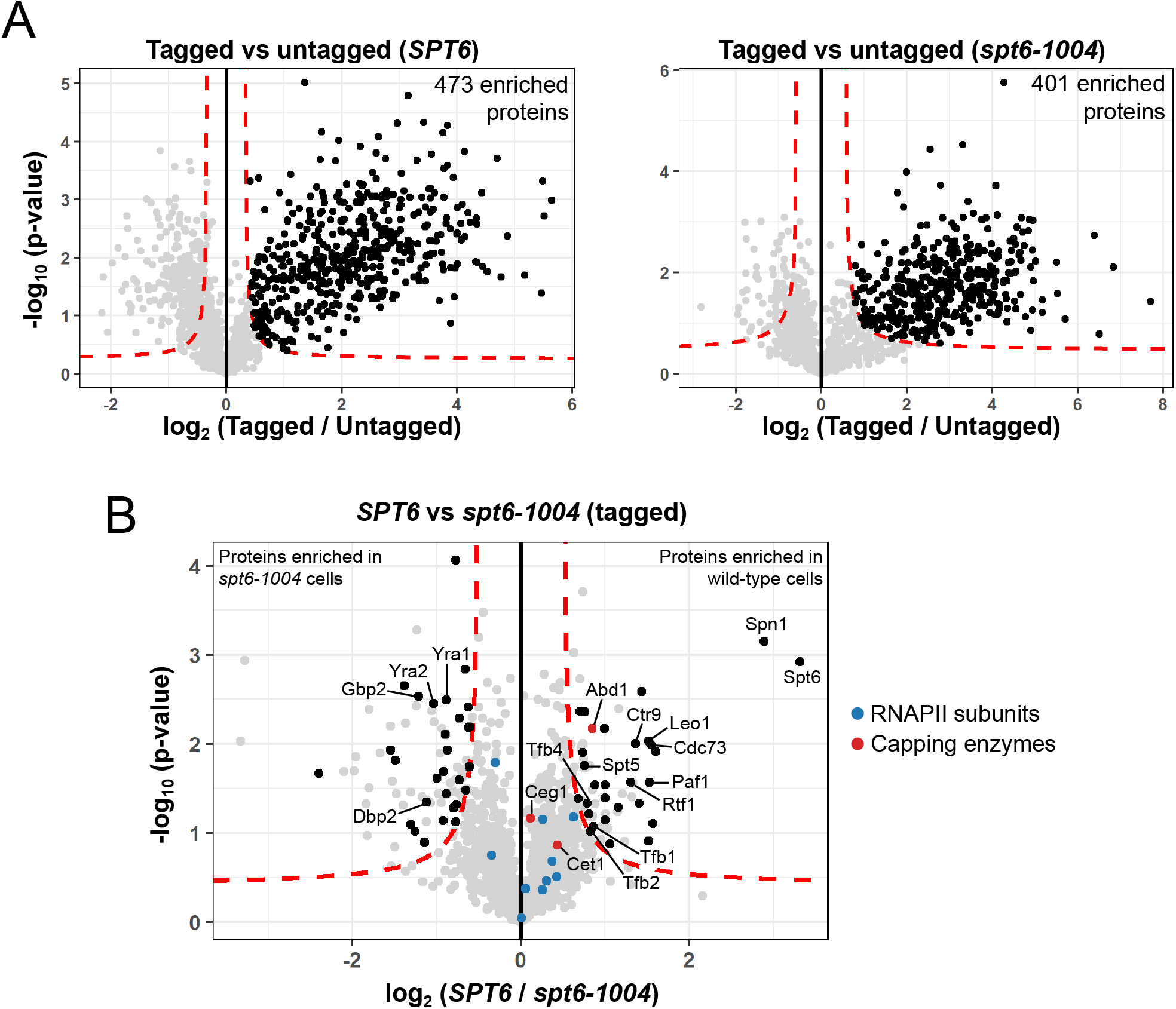
Comparison of RNAPII interacting proteins between wild-type and *spt6-1004* cells. Volcano plot showing comparison of protein abundances between Rpb3-tagged and untagged samples in wild-type and *spt6-1004* cells. Each dot represents a single protein. The dashed red line indicates a permutation based false-discovery rate cutoff of 0.05. Proteins with a positive fold change and above this threshold are considered to be enriched in the tagged sample. (B) Volcano plot showing comparison of protein abundances between wild-type and *spt6-1004* cells in Rpb3-tagged strains. Each dot represents a single protein. The dashed red line indicates a permutation based false-discovery rate cutoff of 0.05. The 58 proteins that were identified to be differentially enriched, as well as enriched over the untagged control in either wild-type or *spt6-1004* cells are highlighted in black. RNAPII subunits and capping enzymes are highlighted in color.

Our results from the mass spectrometry analysis revealed some expected and some previously unknown differences in the RNAPII interactome between wild-type and *spt6-1004* cells. As expected, Spt6 was greatly decreased in *spt6-1004* (Figure 2B) due to the depletion of the mutant Spt6 protein after the shift to 37°C. The protein most decreased after Spt6 in *spt6-1004* was the transcription elongation factor Spn1, which is known to physically interact with Spt6 (33,34). Recent evidence has shown that Spt6 is required for the association of Spn1 with RNAPII (37). We also observed decreased association of all PAF complex subunits with RNAPII in *spt6-1004* (Figure 2B), consistent with previous studies in yeast and *Drosophila*, which showed that Spt6 helps to recruit the PAF complex to actively transcribed regions (17,32). The association of Spt5 with RNAPII was also decreased in *spt6-1004* (Figure 2B). A recent structure of the human transcription elongation complex shows Spt6 directly interacting with Spt5 while binding to RNAPII (14). Interestingly, we also identified the mRNA cap methyltransferase Abd1 as a protein decreased in RNAPII complexes in *spt6-1004*, unlike the two other capping enzymes, Cet1 and Ceg1 (Figure 2B). There were also several proteins apparently enriched in *spt6-1004* cells, including mRNA export and rRNA processing factors (Table 1). In summary, our results suggest that Spt6 regulates the association of a number of transcription factors with RNAPII.

To independently test for differential association of some of these proteins with RNAPII in wild-type and *spt6-1004* cells, we purified Rpb3 from uncrosslinked cells and tested for co-immunoprecipitation of differential interactors. The co-immunoprecipitations were done from cells harvested both before (30°C) and after the temperature shift (37°C). This helped to determine if the altered association of any factor might be primarily due to a specific defect in the *spt6-1004* mutant (30°C) or depletion of Spt6 protein (37°C). We observed that the association of Spt5 with Rpb3 was unchanged in *spt6-1004* cells at 30°C, but decreased at 37°C as compared to wild-type cells (Figure 3A,B). This result agrees with our mass spectrometry data and with previously published data in *Drosophila* cells (17), which shows that the level of Spt5 association with RNAPII is dependent on Spt6 protein levels. A similar observation was also made for Spn1 (Figure 3A,B), although Spn1 protein levels were decreased in *spt6-1004* cells at 37°C, indicating that the reduced association of Spn1 with Rpb3 in *spt6-1004* could be explained in part by reduced Spn1 protein levels. However, a previous study has shown that the association of Spn1 with Rpb3 is dependent on Spt6 (37). The association of the H3K36 methyltransferase Set2 with Rpb3 was also decreased in *spt6-1004* cells at both temperatures (Figure 3A,B). This agrees with our previously published ChIP-seq data (40), where we observed a modest decrease in Set2 recruitment to chromatin in *spt6-1004* cells at 30°C. These co-immunoprecipitation experiments, then, support our mass spectrometry data and have revealed a role for Spt6 regulating the level of Spt5 associated with RNAPII.

**Figure 3.**
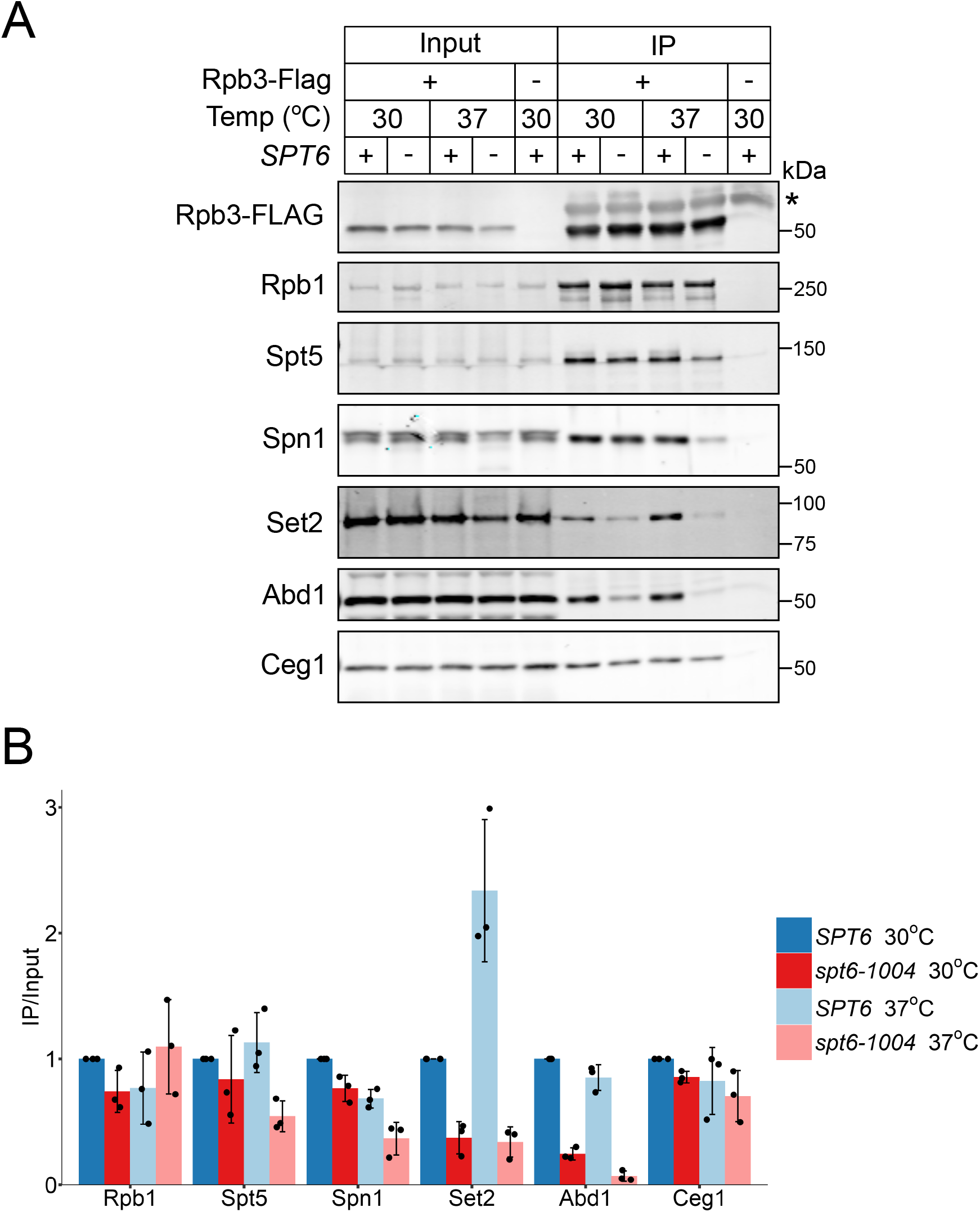
Interactions of transcription elongation factors with Rpb3 in wild-type and *spt6-1004* cells. Co-immunoprecipitation of the indicated proteins with FLAG-tagged Rpb3 in wild-type (+) and *spt6-1004* (-) cells before and after a shift to the non-permissive temperature (37°C). The asterisk (*) indicates the detection of the IgG heavy chain. The images shown here are representative of three independent biological replicates. (B) Quantification of the western blots shown in (A). The immunoprecipitated protein levels were normalized to both input levels of the same protein and immunoprecipiated Rpb3-FLAG levels. The error bars represent mean +/- standard deviation for three replicates.

A novel finding from our mass spectrometry data was the finding that Abd1 is dependent upon Spt6 for association with RNAPII. Our co-immunoprecipitation results also show reduced association of Abd1 with Rpb3 in *spt6-1004* cells, even though global Abd1 protein levels remained unaffected across all conditions tested (Figure 3). Since Spt5 is involved in recruiting Abd1 to chromatin in *S. cerevisiae* (46), the observed differences in Abd1 binding could be due to decreased binding of Spt5 to RNAPII in *spt6-1004*. However, we observed that Abd1 association with Rpb3 was decreased in *spt6-1004* even at 30°C where the association of Spt5 with Rpb3 was unaffected, suggesting that Spt6 regulates the Abd1-RNAPII interaction independently of Spt5. The co-immunoprecipitation of a different capping enzyme, Ceg1, with Rpb3 was unaffected in *spt6-1004* cells (Figure 3), indicating that the altered Rpb3-Abd1 interaction was specific to Abd1 and is not a general property of capping enzymes. Thus, our data reveals a previously unknown role for Spt6 in promoting the association of Abd1 with RNAPII.

We also tested for co-immunoprecipitation of some of the proteins that were enriched in *spt6-1004* in our mass spectrometry data (Dbp2, Chd1, and Iwr1). Unexpectedly, the association of these proteins with Rpb3 was decreased in *spt6-1004*, in contrast to the results from mass spectrometry. Upon further investigation, western blots showed that the levels of some of these proteins were decreased in *spt6-1004* cells following a temperature shift (data not shown). While we cannot yet reconcile our mass spectrometry data with these results, it is possible that, despite loss of total protein levels, a higher proportion of the remaining protein binds to RNAPII transiently, which can be captured only in the presence of the cross-linking that was used in the Bio-TAP XL method.

### Genetic interactions of capping enzyme mutants with *spt6-1004*

Given our observation of decreased association of Abd1 with RNAPII in *spt6-1004*, we reasoned that *abd1* mutations might exacerbate some of the mutant phenotypes observed in *spt6-1004* strains. As *ABD1* is essential for viability, we tested this idea using two *abd1* temperature-sensitive mutations that cause defects in mRNA capping and transcription *in vivo* (47,48). For both, we constructed *abd1 spt6-1004* double mutants by plasmid shuffling (Figure 4A). Our results show that both *abd1-5* and *abd1-8* caused inviability at 30°C when combined with *spt6-1004* (Figure 4B), suggesting that fully functional *ABD1* is required when *SPT6* is mutated. To test if this is a general feature of capping enzyme mutants, we also combined *spt6-1004* with *cet1* and *ceg1* mutations using the same strategy outlined in Figure 4A. The mutations tested were *cet1-401, cet1-438, ceg1-3*, and *ceg1-13*, all of which are temperature sensitive and display defects in mRNA capping (49,50). We observed that *cet1-438* and *cet1-401* are viable when combined with *spt6-1004* (Figure 4C). On the other hand, *ceg1-3 spt6-1004* was inviable, and *ceg1-13 spt6-1004* grew poorly at 30°C (Figure 4D). Our results suggest that there is a functional interaction between Spt6 and mRNA capping, although not specific for Abd1. The greatly increased number of new transcription initiation sites in *spt6-1004* (22) might make the cells hyper-sensitive to impaired capping activity.

**Figure 4.**
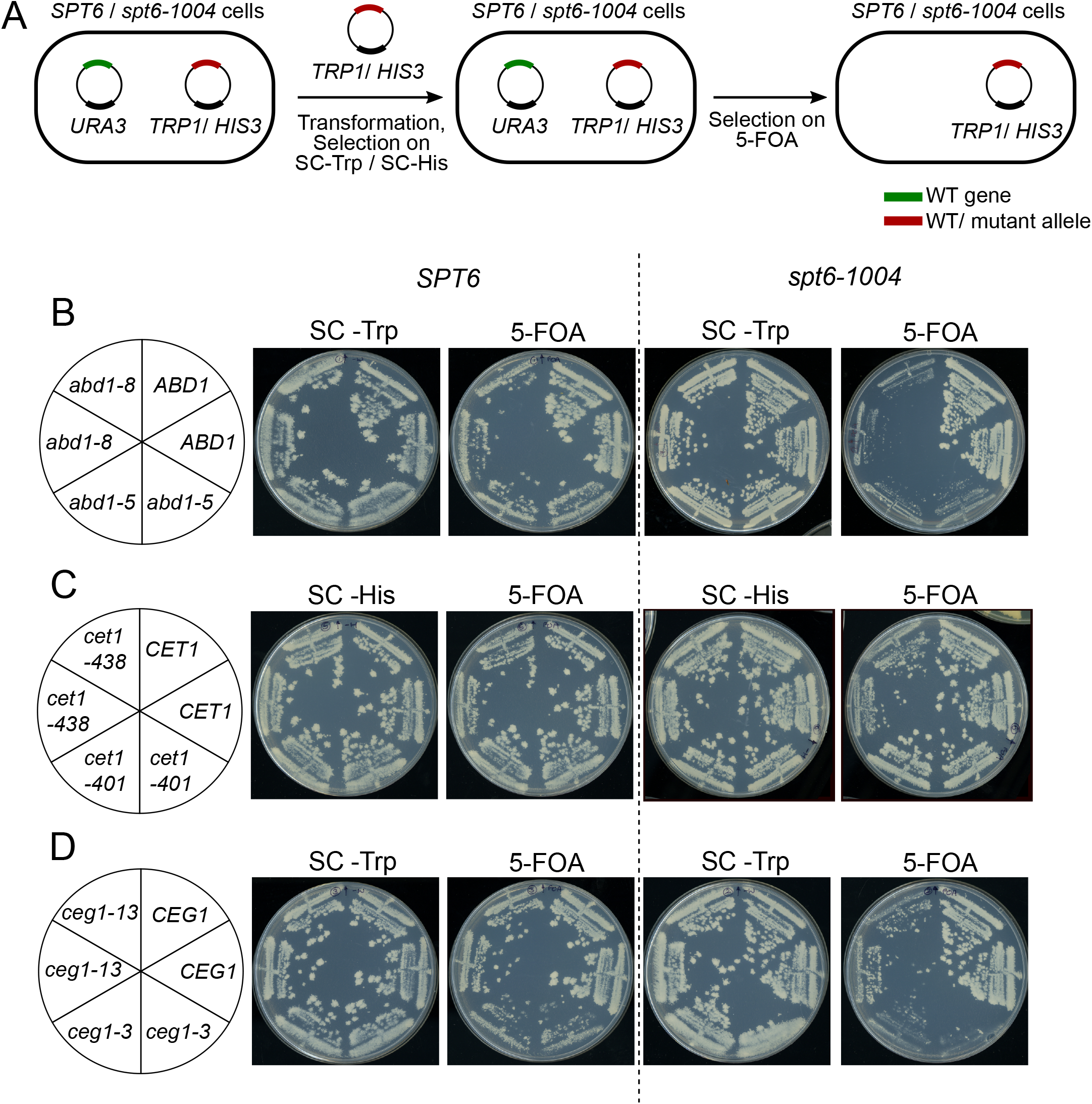
Genetic interactions of capping enzyme mutants with *spt6-1004*. (A) Schematic outlining the plasmid shuffling strategy used to introduce capping enzyme mutants in wild-type and *spt6-1004* cells (B) Growth of wild-type and *spt6-1004* cells expressing the indicated *ABD1* allele. *abd1Δ* or *spt6-1004 abd1Δ* strains expressing wild-type *ABD1* on a *URA3* plasmid were transformed with the indicated *ABD1* allele on a *TRP1* plasmid. Individual transformants were replica plated on SC-Trp or 5-FOA medium and grown for 3 days. (C,D) Similar growth assays were conducted for *CET1* (C) and *CEG1* (D) alleles expressed on *HIS3* and *TRP1* plasmids respectively.

### Genome-wide localization of Abd1 and Spt5 in *spt6-1004*

Previous studies have indicated the requirement of RNAPII and Spt5 for the recruitment of Abd1 during transcription elongation (46,51,52). Our results suggest that Spt6 contributes to the recruitment of Abd1 to chromatin, either directly or possibly via decreased recruitment of RNAPII or Spt5. To test these ideas, we performed ChIP-seq for Abd1-HA, Rpb1, and Spt5-V5 in wild-type and *spt6-1004* cells. For this experiment, cultures were grown at 30°C, the temperature at which we observed defective Rpb3-Abd1 co-immunoprecipitation in *spt6-1004* despite unaltered Spt6 protein levels. To permit quantitative comparison of factor recruitment between different samples, we used chromatin from *S. pombe* for spike-in normalization (Supplementary Figure 2A). Each experiment was done in triplicate and the replicates correlated well with one another (Supplementary Figure 2B).

In wild-type cells, ChIP-seq of Abd1, Spt5, and RNAPII showed the expected patterns of occupancy based on previous studies. Abd1 levels peaked at the 5’ ends of genes at ∼160 bp downstream of the transcription start site, and occupancy at lower levels is observed throughout the gene body (Supplementary Figure 3A), consistent with a previous study (46). Abd1 occupancy showed a sharp decrease following the cleavage and polyadenylation site, suggesting that its association with gene bodies is transcription-dependent. In support of its dependence on transcription, we also observed that Abd1 levels over gene bodies correlated with gene expression levels (Supplementary Figure 3B). The pattern of Spt5 occupancy over gene bodies mirrored that of RNAPII (Supplementary Figure 3B, C, E, F) (16). This is consistent with its association with RNAPII early in transcription and requirement for productive elongation (14,29,53). As expected, the levels of Spt5 and RNAPII occupancy over genes also correlated with their expression level (16) (Supplementary Figure 3E, F).

In an *spt6-1004* mutant, we saw striking differences from wild-type. First, we observed a global decrease in RNAPII occupancy in *spt6-1004*, consistent with previous results (40) (Figure 5C, F). Second, we also observed a global decrease in both Abd1 and Spt5 ChIP-seq levels (Figure 5A, B, D, E). These are likely caused, at least in part, by the decreased levels of RNAPII in *spt6-1004*, as recruitment of both factors to chromatin is dependent on RNAPII (14,51,53). However, although RNAPII levels were decreased uniformly over transcribed regions in *spt6-1004* (Figure 5C), the levels of Abd1 and Spt5 did not follow the same pattern. Abd1 levels showed a greater decrease over the 5’ ends of genes as compared to the decrease over gene bodies (Figure 5A, D). In contrast, the levels of Spt5 were more affected over the gene bodies as compared to the 5’ ends of genes (Figure 5B, E). This suggests that Spt6 is required for maintaining the levels of Abd1 over the 5’ ends of genes and of Spt5, after its recruitment to the transcription elongation complex. Finally, the loss of Abd1, Spt5 and RNAPII occupancy in *spt6-1004* was more prominent at highly expressed genes (Figure 5D-F, Supplementary Figure 4), highlighting the importance of Spt6 during transcription, and suggesting a greater dependence on Spt6 for Abd1 recruitment at highly transcribed genes. Example occupancy levels of Abd1, Spt5, and RNAPII at the highly transcribed *ACT1* gene are shown in Figure 5G. In summary, our results suggest that Spt6 regulates Abd1 and Spt5 recruitment over transcribed regions.

**Figure 5.**
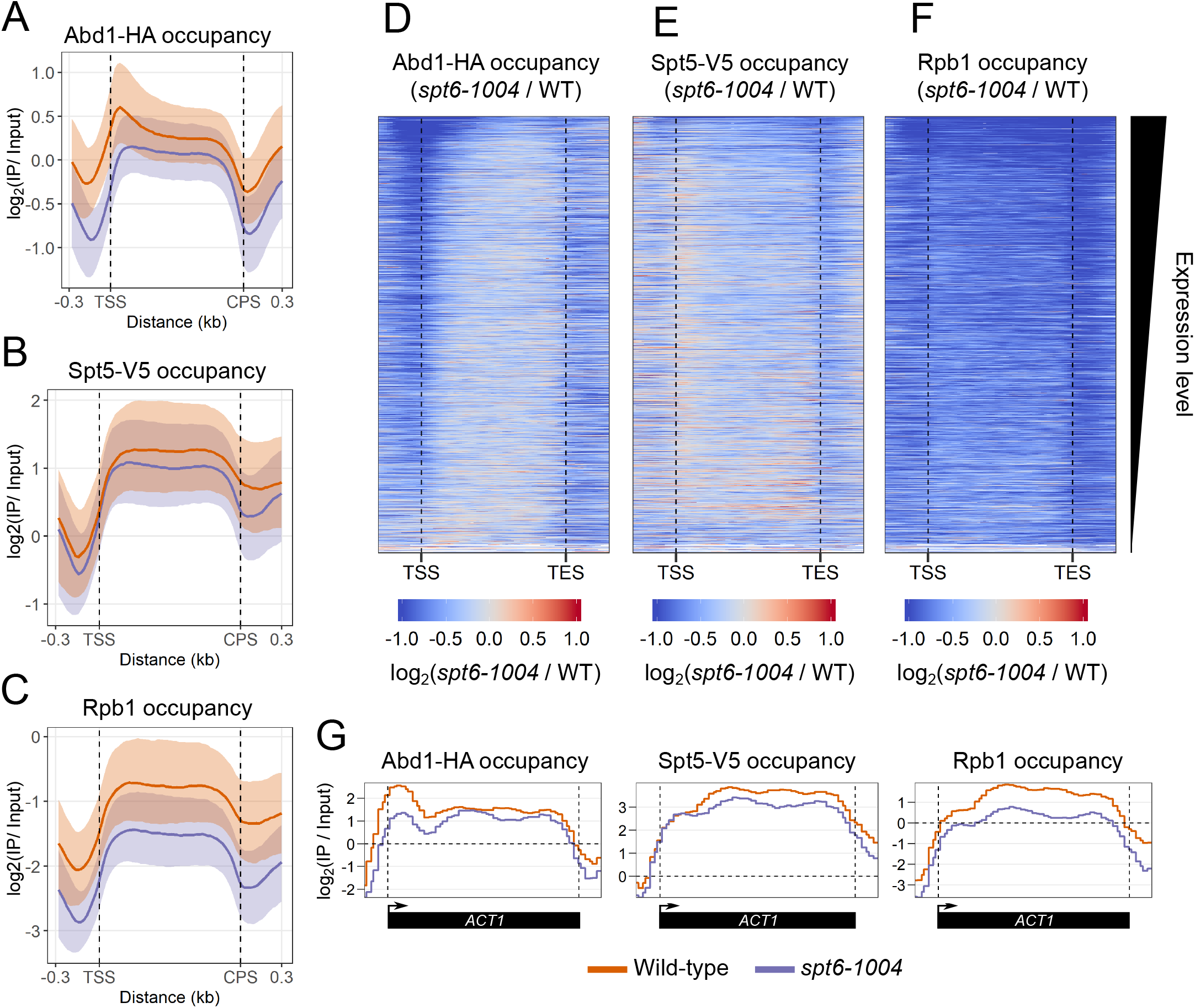
Comparison of genome-wide occupancies of Abd1, Spt5 and RNAPII in wild-type and *spt6-1004*. (A-C) Metagene plots of fold change in Abd1-HA (A), Spt5-V5 (B), and Rpb1 (C) occupancy in wild-type and *spt6-1004* as compared to wild-type cells for 3522 non-overlapping protein-coding genes. The occupancy in each genotype has been normalized to their respective inputs. The trace indicates the median occupancy at a given position. The shaded area represents the interquartile range. (D-F) Heatmaps of fold change in Abd1-HA (D), Spt5-V5 (E), and Rpb1 (F) occupancy in *spt6-1004* as compared to wild-type cells for 3522 non-overlapping protein-coding genes. The occupancy in each genotype has been normalized to their respective inputs. All values that had a log fold change of below -1 or above 1 are set to -1 and 1 respectively. The genes are arranged in decreasing order of expression level as determined from RNA-Seq of wild-type cells (Ref. 37). (G) Representative traces of Abd1-HA, Spt5-V5 and Rpb1 occupancy in wild-type and *spt6-1004* cells at the highly transcribed *ACT1* gene.

## Discussion

We have identified a role for Spt6 in maintaining the association of transcription elongation factors with RNAPII. The RNAPII-interacting proteins that we have identified in wild-type cells are similar to those identified in previous studies (54,55), indicating that BioTAP-XL can be used to successfully purify protein complexes in yeast. Of the RNAPII-interacting proteins detected, we identified 58 that were specifically depleted or enriched in RNAPII complexes in an *spt6-1004* mutant. These included factors previously known to interact with Spt6 (such as Spt5 and Spn1) as well as factors whose relationship with Spt6 was previously unknown, most notably Abd1. Our results suggest that the altered accumulation of some of these factors is due to their reduced protein levels in *spt6-1004*, which may either be due to changes in gene expression or protein stability. We followed up on one of the proteins whose steady-state level remained unchanged in *spt6-1004* – the mRNA cap methyltransferase Abd1 and showed that the occupancy of Abd1 on chromatin in *spt6-1004* is decreased genome-wide. We also identified a decreased occupancy for Spt5 genome-wide, revealing another requirement for Spt6 in regulating transcriptional processes.

We showed that the levels of Abd1 are most decreased at the 5’ ends of genes in the *spt6-1004* mutant. Two plausible models can explain this observation. In the first model, Spt6 plays a role in recruiting Abd1 specifically at the 5’ end. While Abd1 occupancy depends on Spt5 throughout the gene, it may additionally depend on Spt6 levels towards the 5’ ends of genes. This would explain why we see a greater defect in Abd1 recruitment at the 5’ end even when the Spt5 recruitment defect is lowest in this region. A second possibility is that levels of Ser-5 phosphorylated CTD of Rpb1, which is required for normal recruitment of Abd1 (51), might be decreased in *spt6-1004*. In support of this hypothesis, in the *spt6-1004* mutant we also observe decreased RNAPII association of Tfb1, Tfb2 and Tfb4 (Figure 2B), which are components of the TFIIH complex that contains the Kin28 kinase that phosphorylates the Ser-5 residue of the CTD. Hence, although required for Abd1 recruitment, the role of Spt6 in this process might be through an indirect effect on transcription.

By ChIP-Seq, we also observed that RNAPII and Spt5 occupancies are decreased over transcribed regions genome-wide in *spt6-1004*. Our previous data also showed a global decrease in RNAPII ChIP-seq signal in *spt6-1004* (40), although to a lesser extent. The higher magnitude observed here may be due to the presence of the V5 tag on Spt5 in the same strain, leading to genetic interactions between *SPT5* and *spt6-1004* that impair transcription. However, we do not observe any effect of *SPT5-V5* on the temperature sensitivity or Spt^-^ phenotypes of *spt6-1004*. While RNAPII levels show a uniform decrease over gene bodies, the levels of Spt5 decrease more at the 3’ ends of genes as compared to the 5’ ends, indicating that Spt6 might be required for continued association of Spt5 with RNAPII during transcription elongation. This agrees with previously published data showing reduced recruitment of Spt5 to the *Hsp70* gene body upon knockdown of *SPT6* in *Drosophila* cells (17). This is also in line with the structure of the human transcription elongation complex, where part of Spt5 is sandwiched between RNAPII and Spt6 (14). Loss of Spt6 might allow dissociation of Spt5, leading to the lower levels of RNAPII observed genome-wide in *spt6-1004*.

The decreased association of Abd1 over transcribed regions in *spt6-1004* likely has functional consequences. This is supported by our genetic data showing loss of viability upon combining *abd1* mutations with *spt6-1004*. First, it can affect mRNA capping, possibly leading to the production of transcripts that have non-methylated mRNA caps. Such incompletely capped mRNAs are observed even in wild-type cells under stress conditions and the exonucleases Rai1 and Dxo1 are involved in the recognition and degradation of these transcripts (56-58). This indicates that inefficient mRNA capping can occur under physiological conditions and that the cell has developed quality control mechanisms to prevent the accumulation of aberrantly capped transcripts. Second, the decreased association of Abd1 over gene bodies can affect transcription. Abd1 has been previously observed to positively promote transcription in yeast as well as in humans (47,59). Future experiments will help determine if the reduced interaction of Abd1 with RNAPII may be contributing to some of the phenotypes observed in *spt6-1004*.

## Experimental procedures

### Strains and media

All *S. cerevisiae* and *S. pombe* strains used are listed in Table S1. All plasmids used are listed in Table S2. All oligos used for strains constructions are listed in Table S3. All *S. cerevisiae* strains were grown in YPD medium (1% yeast extract, 2% peptone and 2% glucose) unless mentioned otherwise. All *S. pombe* strains were grown in YES medium (0.5% yeast extract, 3% glucose, 225 mg/l each of adenine, histidine, leucine, uracil and lysine). For experiments involving temperature shifts, cells were first grown to OD_600_ ≈ 0.6 (∼2⨯10^7^ cells/ml) at 30°C. One volume of culture was mixed with an equal volume of YPD at 42°C. The cells were then grown at 37°C for 80 minutes. All strains were constructed by standard yeast crosses or transformations (60). For tagging Rpb3 with the BioTAP tag, the DNA sequence encoding the tag was amplified from plasmid FB2729 and the PCR fragment was used to transform yeast strain FY57, resulting in integration into the *S. cerevisiae* genome, replacing the stop codon in the *RPB3* gene. Tagged *RPB3* was introduced in *spt6-1004* through a genetic cross with RGC42. For tagging *ABD1* with 3xHA, the DNA encoding the tag was amplified from the plasmid pFA6a-3xHA-kanMX6 (61), and the PCR fragment was transformed into a wild-type yeast strain, resulting in integration into the *S. cerevisiae* genome, replacing the stop codon in the *ABD1* gene. *SPT5-3xV5* and *spt6-1004* were introduced in the tagged *ABD1* strain through standard genetic crosses. Wild-type or mutated versions of genes encoding capping enzymes were introduced into yeast strains by plasmid shuffling, using plasmids CE113, CE333, and CE339 for *CET1*; RGBO5, RGBO6, and RGBO7 for *CEG1*; and RGBO8, RGBO9, and RGBO10 for *ABD1*.

### Purification of RNAPII complexes by BioTAP-XL and analysis by mass spectrometry

BioTAP-XL was done as previously described (42), with modifications to adapt the protocol from *D. melanogaster* cells to *S. cerevisiae* cells. Yeast strains RGC11, RGC14, FY57, and RGC42 were grown in 1.5 L of YPD supplemented with 6 µM biotin at 30°C to an OD_600_≈0.6. This was followed by addition of 1.5 L of YPD (supplemented with 6 µM biotin) at 42°C and growth at 37°C for 80 minutes. The yeast cultures were then rapidly cooled by transferring the cells to a flask with 0.3x volume of media at 4°C, and formaldehyde was immediately added to a final concentration of 1%. The cultures were incubated with shaking at room temperature for 30 minutes. Glycine was then added to a final concentration of 125 mM and the incubation was continued for 5 minutes. The cells were collected by filtration onto a 0.45 µm nitrocellulose membrane filter. The cells were then scraped off the filter, inserted into a syringe, and expelled into liquid nitrogen. The frozen cells were lysed in a mixer mill for 6 cycles, 3 minutes each at 15 Hz, with incubation in liquid nitrogen between each cycle, to form a “grindate.” The grindate was suspended in 12 ml of cold FA lysis buffer (50 mM HEPES-KOH pH 7.5, 140 mM NaCl, 1mM EDTA, 0.1% sodium deoxycholate, 0.1% Triton X-100, 0.05% SDS, 2 µg/ml leupeptin, 2 µg/ml pepstain, 1 mM PMSF) and divided equally between 15 tubes. The sample in each tube was pelleted by centrifugation, resuspended in 640 µl of FA lysis buffer, and sonicated in a Qsonica machine for 25 minutes (30 seconds on, 30 seconds off, 70% amplitude). The sonicated samples were centrifuged at 12,500 rpm for 30 minutes at 4°C. The supernatants from individual tubes were pooled and incubated with 1.8 ml of IgG beads (pre-washed with FA lysis buffer) overnight at 4°C.

Following the IP, the beads were washed three times with RIPA buffer (10 mM Tris-HCl pH 8.0, 140 mM NaCl, 1mM EDTA, 1% Triton X-100, 0.1% SDS). For each wash, the resuspended beads were incubated at 4°C for 10 minutes with end-over-end rotation. The beads were then washed once with TEN140 buffer (10 mM Tris-HCl pH 8.0, 140 mM NaCl, 1mM EDTA) with end-over-end rotation for 2 minutes at room temperature. The protein complexes were eluted twice by incubation with 20 ml of freshly made IgG elution buffer (100 mM Tris-HCl pH 8.0, 200 mM NaCl, 6M Urea, 0.2% SDS) with end-over-end rotation for one hour at room temperature. The eluted sample was concentrated in an Amicon Ultra-15 column (10kDa cutoff, Millipore) and buffer exchange was done four times with 12 ml of TEN140 buffer. The resulting 1 ml of concentrated sample was brought up to 2.8 ml with RIPA buffer and incubated with 1 ml of streptavidin agarose beads (pre-washed with TEN140 buffer).

Following the affinity pulldown, the beads were washed once with TEN140 buffer with 0.1% Triton-X 100, twice with IgG elution buffer, twice with IgG elution buffer without SDS, and once with TEN140 buffer for 5 minutes each at 4°C. The beads were then washed seven times with 50 mM ammonium bicarbonate for 5 minutes each at room temperature and then suspended in 800 µl of 50 mM ammonium bicarbonate and split evenly between two tubes. Ten µl of trypsin was added to one tube, which was then incubated overnight at 37°C with end-over-end rotation. The beads in the other tube were boiled for 25 minutes in reverse cross-linking buffer (250 mM Tris-HCl pH 8.8, 2% SDS, 0.5 M β-mercaptoethanol) and the resulting supernatant was stored at -70°C.

One µl of 100% formic acid was added to the trypsinization reaction. The beads were centrifuged at 3000 rpm for 5 minutes at 25°C and the supernatant containing the digested peptides was transferred to a new tube. The beads were washed three times with a solution of 25% acetonitrile and 0.1% formic acid, and the supernatants after each wash was pooled with the initial supernatant. A C18 spin tip was equilibrated with 50 µl of 100% acetonitrile and twice with 50 µl of 0.1% tri-fluoroacetic acid. The peptides were passed twice through the C18 spin tip, which was then washed twice with 0.1% tri-fluoroacetic acid. The peptides were then eluted from the resin once with 30 µl of 50% acetonitrile and once with 30 µl of 100% acetonitrile. The eluted peptides were dried completely in a SpeedVac vacuum concentrator and the dried peptides were stored at -70°C until analysis by mass spectrometry.

Mass spectrometry analysis was conducted by Ryan Kunz at the Taplin Mass Spectrometry facility at Harvard Medical School. The peptides from each sample were labeled with tandem mass tags and quantification of peptide abundances among the different samples was done at the MS3 stage (44).

### Analysis of mass spectrometry data

Following reporter ion quantification, the peptide intensities for each protein were summed and normalized to the total peptide intensity in each sample. The data were processed using Perseus software (45). The summed peptide intensities for each protein were log transformed and missing values (for proteins that had summed peptide intensity = 0) were imputed from a normal distribution. The samples were grouped by genotype and a t-test was conducted to compare the signal for each protein between two samples of interest. A permutation-based false discovery rate threshold was calculated and used to identify proteins as being significantly enriched in one sample versus the other. Volcano plots were generated using custom R scripts.

### Co-immunoprecipitation assays

For each co-immunoprecipitation experiment, 50 ml of yeast cells at OD_600_≈0.6 (∼2⨯10^7^ cells/ml) were harvested. The cell pellets were suspended in 500 µl of IP buffer (20 mM HEPES-KOH, pH 7.6, 125 mM potassium acetate, 1 mM EDTA, 20% glycerol, 1mM DTT, 1% NP-40, 1x Sigma protease inhibitor cocktail). One ml of acid-washed glass beads was added to each tube and the cells were lysed by bead beating for 8 minutes with incubation on ice for 3 minutes after each minute. The resulting lysate was centrifuged at 12,500 rpm for 10 minutes at 4°C. The supernatant was transferred to a new tube. Protein concentrations in the whole cell lysates were measured by Bradford assay (62). One mg of protein in a final volume of 500 µl was incubated with 20 µl of anti-flag agarose beads (Sigma) pre-washed with IP buffer. The IP was carried out for 2 hours at 4°C. The beads were then washed thrice with 500 µl of IP buffer following which the beads were boiled for 5 minutes in 50 µl of Modified SDS buffer (60 mM Tris-HCl pH 6.8, 4% β-mercaptoethanol, 4% SDS, 0.01% bromophenol blue and 20% glycerol). The beads were centrifuged at 12,500 rpm for 1 minute at room temperature and 15 µl of the supernatant was loaded on sodium dodecyl sulphate-polyacrylamide gel electrophoresis (SDS-PAGE) gels for western blotting.

### Western blotting and antibodies

Whole cell extracts from yeast were prepared as described previously (40). Primary antibodies used for western blotting were anti-Set2 (1:8000, provided by Brian Strahl), anti-Abd1 (1:1000, provided by Stephen Buratowski), anti-Ceg1 (1:2000, provided by Stephen Buratowski), anti-Spn1 (1:8000, provided by Laurie Stargell), anti-Spt6 (1:10,000, provided by Brian Strahl), anti-Rpb1 (1:1000, Millipore, 8WG16), anti-HA (1:5000, Abcam, ab9110), anti-Flag (1:5000, Sigma, F3165), anti-V5 (1:5000, Invitrogen, R960-25), anti-Pgk1 (1:10,000, Life Technologies 459250) and anti-Act1 (1:10,000, Abcam, ab8224). Secondary antibodies used were: goat anti-rabbit IgG (1:10,000, Licor IRDye 680RD) and goat anti-mouse IgG (1:20,000, Licor, 800CW). Quantification of western blots was done using Licor ImageStudio software.

### Chromatin immunoprecipitation

ChIP-qPCR and ChIP-seq library preparation was done as described previously (40), with minor modifications. Each sample was spiked-in with *S. pombe* chromatin from strains FWP566 and FWP485 to a final concentration of 7.5% for each strain prior to the immunoprecipitation step. 5µl of anti-HA antibody (Abcam, ab9110) per 500µg of chromatin, 7.5 µl of anti-V5 antibody (Invitrogen, R960-25) per 500µg of chromatin and 10 µl of anti-Rpb1 antibody (Millipore, 8WG16) per 500 µg of chromatin were used for immunoprecipitation of Abd1-HA, Spt5-V5 and Rpb1 respectively. Oligonucleotides used for ChIP-qPCR are listed in Table S3.

### ChIP-Seq data analysis

De-multiplexing, alignment, spike-in normalization and generation of coverage files from FASTQ files was done as described previously (40). Commands from the deeptools suite (63) was used for normalizing libraries to each other and generating matrices suitable for plotting heatmaps and metagenes. Metagenes and heatmaps were generated using custom R scripts.

## Data availability

The RNA-seq and ChIP-seq data sets are available in the GEO repository, accession number GSE171953. They can be accessed at https://www.ncbi.nlm.nih.gov/geo/query/acc.cgi?acc=GSE171953. All other relevant data supporting the key findings of this study are available within this article and its Supplementary Data.

## Supporting information

This article contains supporting information.

## Acknowledgments

We thank Ryan Kunz for advice and help with the mass spectrometry experiments and analysis, Beate Schwer for providing us with the *ABD1* and *CEG1* plasmids, and Steve Buratowski for providing us with the *CET1* plasmids and helpful discussions. We also thank Catherine Weiner and James Warner for helpful comments on the manuscript.

## Author contributions

R.G. and F.W. designed the experiments. R.G. carried out all of the experiments. R.G. and F.W. wrote the paper.

## Funding and additional information

This work was supported by NIH grants R01GM32967 and R01GM120038 to F.W.

## References

1. Zhou, Q., Li, T., and Price, D. H. (2012) RNA polymerase II elongation control. Annu Rev Biochem 81, 119–143

2. Venkatesh, S., and Workman, J. L. (2015) Histone exchange, chromatin structure and the regulation of transcription. Nat Rev Mol Cell Biol 16, 178–189

3. Swanson, M. S., Carlson, M., and Winston, F. (1990) SPT6, an essential gene that affects transcription in Saccharomyces cerevisiae, encodes a nuclear protein with an extremely acidic amino terminus. Molecular and cellular biology 10, 4935–4941

4. Duina, A. A. (2011) Histone Chaperones Spt6 and FACT: Similarities and Differences in Modes of Action at Transcribed Genes. Genet Res Int 2011, 625210

5. Kok, F. O., Oster, E., Mentzer, L., Hsieh, J. C., Henry, C. A., and Sirotkin, H. I. (2007) The role of the SPT6 chromatin remodeling factor in zebrafish embryogenesis. Dev Biol 307, 214–226

6. Bourbon, H. M., Gonzy-Treboul, G., Peronnet, F., Alin, M. F., Ardourel, C., Benassayag, C., Cribbs, D., Deutsch, J., Ferrer, P., Haenlin, M., Lepesant, J. A., Noselli, S., and Vincent, A. (2002) A P-insertion screen identifying novel X-linked essential genes in Drosophila. Mech Dev 110, 71–83

7. Meyers, R. M., Bryan, J. G., McFarland, J. M., Weir, B. A., Sizemore, A. E., Xu, H., Dharia, N. V., Montgomery, P. G., Cowley, G. S., Pantel, S., Goodale, A., Lee, Y., Ali, L. D., Jiang, G., Lubonja, R., Harrington, W. F., Strickland, M., Wu, T., Hawes, D. C., Zhivich, V. A., Wyatt, M. R., Kalani, Z., Chang, J. J., Okamoto, M., Stegmaier, K., Golub, T. R., Boehm, J. S., Vazquez, F., Root, D. E., Hahn, W. C., and Tsherniak, A. (2017) Computational correction of copy number effect improves specificity of CRISPR-Cas9 essentiality screens in cancer cells. Nature genetics 49, 1779–1784

8. Liu, J., Zhang, J., Gong, Q., Xiong, P., Huang, H., Wu, B., Lu, G., Wu, J., and Shi, Y. (2011) Solution structure of tandem SH2 domains from Spt6 protein and their binding to the phosphorylated RNA polymerase II C-terminal domain. The Journal of biological chemistry 286, 29218–29226

9. Dengl, S., Mayer, A., Sun, M., and Cramer, P. (2009) Structure and in vivo requirement of the yeast Spt6 SH2 domain. Journal of molecular biology 389, 211–225

10. Diebold, M. L., Loeliger, E., Koch, M., Winston, F., Cavarelli, J., and Romier, C. (2010) Noncanonical tandem SH2 enables interaction of elongation factor Spt6 with RNA polymerase II. The Journal of biological chemistry 285, 38389–38398

11. Sdano, M. A., Fulcher, J. M., Palani, S., Chandrasekharan, M. B., Parnell, T. J., Whitby, F. G., Formosa, T., and Hill, C. P. (2017) A novel SH2 recognition mechanism recruits Spt6 to the doubly phosphorylated RNA polymerase II linker at sites of transcription. eLife 6

12. Close, D., Johnson, S. J., Sdano, M. A., McDonald, S. M., Robinson, H., Formosa, T., and Hill, C. P. (2011) Crystal structures of the S. cerevisiae Spt6 core and C-terminal tandem SH2 domain. Journal of molecular biology 408, 697–713

13. Sun, M., Lariviere, L., Dengl, S., Mayer, A., and Cramer, P. (2010) A tandem SH2 domain in transcription elongation factor Spt6 binds the phosphorylated RNA polymerase II C-terminal repeat domain (CTD). The Journal of biological chemistry 285, 41597–41603

14. Vos, S. M., Farnung, L., Boehning, M., Wigge, C., Linden, A., Urlaub, H., and Cramer, P. (2018) Structure of activated transcription complex Pol II-DSIF-PAF-SPT6. Nature 560, 607–612

15. Ivanovska, I., Jacques, P. E., Rando, O. J., Robert, F., and Winston, F. (2011) Control of chromatin structure by spt6: different consequences in coding and regulatory regions. Molecular and cellular biology 31, 531–541

16. Mayer, A., Lidschreiber, M., Siebert, M., Leike, K., Soding, J., and Cramer, P. (2010) Uniform transitions of the general RNA polymerase II transcription complex. Nature structural & molecular biology 17, 1272–1278

17. Ardehali, M. B., Yao, J., Adelman, K., Fuda, N. J., Petesch, S. J., Webb, W. W., and Lis, J. T. (2009) Spt6 enhances the elongation rate of RNA polymerase II in vivo. The EMBO journal 28, 1067–1077

18. Endoh, M., Zhu, W., Hasegawa, J., Watanabe, H., Kim, D. K., Aida, M., Inukai, N., Narita, T., Yamada, T., Furuya, A., Sato, H., Yamaguchi, Y., Mandal, S. S., Reinberg, D., Wada, T., and Handa, H. (2004) Human Spt6 stimulates transcription elongation by RNA polymerase II in vitro. Molecular and cellular biology 24, 3324–3336

19. McCullough, L., Connell, Z., Petersen, C., and Formosa, T. (2015) The Abundant Histone Chaperones Spt6 and FACT Collaborate to Assemble, Inspect, and Maintain Chromatin Structure in Saccharomyces cerevisiae. Genetics 201, 1031–1045

20. Bortvin, A., and Winston, F. (1996) Evidence that Spt6p controls chromatin structure by a direct interaction with histones. Science 272, 1473–1476

21. van Bakel, H., Tsui, K., Gebbia, M., Mnaimneh, S., Hughes, T. R., and Nislow, C. (2013) A compendium of nucleosome and transcript profiles reveals determinants of chromatin architecture and transcription. PLoS genetics 9, e1003479

22. Doris, S. M., Chuang, J., Viktorovskaya, O., Murawska, M., Spatt, D., Churchman, L. S., and Winston, F. (2018) Spt6 Is Required for the Fidelity of Promoter Selection. Molecular cell 72, 687-699 e686

23. DeGennaro, C. M., Alver, B. H., Marguerat, S., Stepanova, E., Davis, C. P., Bahler, J., Park, P. J., and Winston, F. (2013) Spt6 regulates intragenic and antisense transcription, nucleosome positioning, and histone modifications genome-wide in fission yeast. Mol Cell Biol 33, 4779–4792

24. Perales, R., Erickson, B., Zhang, L., Kim, H., Valiquett, E., and Bentley, D. (2013) Gene promoters dictate histone occupancy within genes. The EMBO journal 32, 2645–2656

25. Kaplan, C. D., Laprade, L., and Winston, F. (2003) Transcription elongation factors repress transcription initiation from cryptic sites. Science 301, 1096–1099

26. Cheung, V., Chua, G., Batada, N. N., Landry, C. R., Michnick, S. W., Hughes, T. R., and Winston, F. (2008) Chromatin-and transcription-related factors repress transcription from within coding regions throughout the Saccharomyces cerevisiae genome. PLoS biology 6, e277

27. Uwimana, N., Collin, P., Jeronimo, C., Haibe-Kains, B., and Robert, F. (2017) Bidirectional terminators in Saccharomyces cerevisiae prevent cryptic transcription from invading neighboring genes. Nucleic acids research 45, 6417–6426

28. Hartzog, G. A., Wada, T., Handa, H., and Winston, F. (1998) Evidence that Spt4, Spt5, and Spt6 control transcription elongation by RNA polymerase II in Saccharomyces cerevisiae. Genes & development 12, 357–369

29. Endoh, M., Zhu, W., Hasegawa, J., Watanabe, H., Kim, D. K., Aida, M., Inukai, N., Narita, T., Yamada, T., Furuya, A., Sato, H., Yamaguchi, Y., Mandal, S. S., Reinberg, D., Wada, T., and Handa, H. (2004) Human Spt6 Stimulates Transcription Elongation by RNA Polymerase II In Vitro. Molecular and Cellular Biology 24, 3324–3336

30. Swanson, M. S., and Winston, F. (1992) SPT4, SPT5 and SPT6 interactions: effects on transcription and viability in Saccharomyces cerevisiae. Genetics 132, 325–336

31. Van Oss, S. B., Cucinotta, C. E., and Arndt, K. M. (2017) Emerging Insights into the Roles of the Paf1 Complex in Gene Regulation. Trends in biochemical sciences 42, 788–798

32. Kaplan, C. D., Holland, M. J., and Winston, F. (2005) Interaction between transcription elongation factors and mRNA 3’-end formation at the Saccharomyces cerevisiae GAL10-GAL7 locus. The Journal of biological chemistry 280, 913–922

33. Diebold, M. L., Koch, M., Loeliger, E., Cura, V., Winston, F., Cavarelli, J., and Romier, C. (2010) The structure of an Iws1/Spt6 complex reveals an interaction domain conserved in TFIIS, Elongin A and Med26. The EMBO journal 29, 3979–3991

34. McDonald, S. M., Close, D., Xin, H., Formosa, T., and Hill, C. P. (2010) Structure and biological importance of the Spn1-Spt6 interaction, and its regulatory role in nucleosome binding. Molecular cell 40, 725–735

35. Yoh, S. M., Lucas, J. S., and Jones, K. A. (2008) The Iws1:Spt6:CTD complex controls cotranscriptional mRNA biosynthesis and HYPB/Setd2-mediated histone H3K36 methylation. Genes & development 22, 3422–3434

36. Oqani, R. K., Lin, T., Lee, J. E., Kang, J. W., Shin, H. Y., and Il Jin, D. (2019) Iws1 and Spt6 Regulate Trimethylation of Histone H3 on Lysine 36 through Akt Signaling and are Essential for Mouse Embryonic Genome Activation. Scientific reports 9, 3831

37. Reim, N. I., Chuang, J., Jain, D., Alver, B. H., Park, P. J., and Winston, F. (2020) The conserved elongation factor Spn1 is required for normal transcription, histone modifications, and splicing in Saccharomyces cerevisiae. Nucleic acids research 48, 10241–10258

38. Youdell, M. L., Kizer, K. O., Kisseleva-Romanova, E., Fuchs, S. M., Duro, E., Strahl, B. D., and Mellor, J. (2008) Roles for Ctk1 and Spt6 in regulating the different methylation states of histone H3 lysine 36. Molecular and cellular biology 28, 4915–4926

39. Chu, Y., Sutton, A., Sternglanz, R., and Prelich, G. (2006) The BUR1 cyclin-dependent protein kinase is required for the normal pattern of histone methylation by SET2. Molecular and cellular biology 26, 3029–3038

40. Gopalakrishnan, R., Marr, S. K., Kingston, R. E., and Winston, F. (2019) A conserved genetic interaction between Spt6 and Set2 regulates H3K36 methylation. Nucleic acids research

41. Yoh, S. M., Cho, H., Pickle, L., Evans, R. M., and Jones, K. A. (2007) The Spt6 SH2 domain binds Ser2-P RNAPII to direct Iws1-dependent mRNA splicing and export. Genes & development 21, 160–174

42. Alekseyenko, A. A., McElroy, K. A., Kang, H., Zee, B. M., Kharchenko, P. V., and Kuroda, M. I. (2015) BioTAP-XL: Cross-linking/Tandem Affinity Purification to Study DNA Targets, RNA, and Protein Components of Chromatin-Associated Complexes. Curr Protoc Mol Biol 109, 21 30 21–32

43. Thompson, A., Schafer, J., Kuhn, K., Kienle, S., Schwarz, J., Schmidt, G., Neumann, T., Johnstone, R., Mohammed, A. K., and Hamon, C. (2003) Tandem mass tags: a novel quantification strategy for comparative analysis of complex protein mixtures by MS/MS. Anal Chem 75, 1895–1904

44. Ting, L., Rad, R., Gygi, S. P., and Haas, W. (2011) MS3 eliminates ratio distortion in isobaric multiplexed quantitative proteomics. Nat Methods 8, 937–940

45. Tyanova, S., Temu, T., Sinitcyn, P., Carlson, A., Hein, M. Y., Geiger, T., Mann, M., and Cox, J. (2016) The Perseus computational platform for comprehensive analysis of (prote)omics data. Nat Methods 13, 731–740

46. Lidschreiber, M., Leike, K., and Cramer, P. (2013) Cap completion and C-terminal repeat domain kinase recruitment underlie the initiation-elongation transition of RNA polymerase II. Mol Cell Biol 33, 3805–3816

47. Schroeder, S. C., Zorio, D. A., Schwer, B., Shuman, S., and Bentley, D. (2004) A function of yeast mRNA cap methyltransferase, Abd1, in transcription by RNA polymerase II. Mol Cell 13, 377–387

48. Schwer, B., Saha, N., Mao, X., Chen, H. W., and Shuman, S. (2000) Structure-function analysis of yeast mRNA cap methyltransferase and high-copy suppression of conditional mutants by AdoMet synthase and the ubiquitin conjugating enzyme Cdc34p. Genetics 155, 1561–1576

49. Takase, Y., Takagi, T., Komarnitsky, P. B., and Buratowski, S. (2000) The essential interaction between yeast mRNA capping enzyme subunits is not required for triphosphatase function in vivo. Mol Cell Biol 20, 9307–9316

50. Schwer, B., Mao, X., and Shuman, S. (1998) Accelerated mRNA decay in conditional mutants of yeast mRNA capping enzyme. Nucleic Acids Res 26, 2050–2057

51. Schroeder, S. C., Schwer, B., Shuman, S., and Bentley, D. (2000) Dynamic association of capping enzymes with transcribing RNA polymerase II. Genes Dev 14, 2435–2440

52. Pei, Y., and Shuman, S. (2002) Interactions between fission yeast mRNA capping enzymes and elongation factor Spt5. J Biol Chem 277, 19639–19648

53. Andrulis, E. D., Guzman, E., Doring, P., Werner, J., and Lis, J. T. (2000) High-resolution localization of Drosophila Spt5 and Spt6 at heat shock genes in vivo: roles in promoter proximal pausing and transcription elongation. Genes & development 14, 2635–2649

54. Mosley, A. L., Hunter, G. O., Sardiu, M. E., Smolle, M., Workman, J. L., Florens, L., and Washburn, M. P. (2013) Quantitative proteomics demonstrates that the RNA polymerase II subunits Rpb4 and Rpb7 dissociate during transcriptional elongation. Mol Cell Proteomics 12, 1530–1538

55. Tardiff, D. F., Abruzzi, K. C., and Rosbash, M. (2007) Protein characterization of Saccharomyces cerevisiae RNA polymerase II after in vivo cross-linking. Proc Natl Acad Sci U S A 104, 19948–19953

56. Chang, J. H., Jiao, X., Chiba, K., Oh, C., Martin, C. E., Kiledjian, M., and Tong, L. (2012) Dxo1 is a new type of eukaryotic enzyme with both decapping and 5’-3’ exoribonuclease activity. Nat Struct Mol Biol 19, 1011–1017

57. Jiao, X., Chang, J. H., Kilic, T., Tong, L., and Kiledjian, M. (2013) A mammalian pre-mRNA 5’ end capping quality control mechanism and an unexpected link of capping to pre-mRNA processing. Mol Cell 50, 104–115

58. Jiao, X., Xiang, S., Oh, C., Martin, C. E., Tong, L., and Kiledjian, M. (2010) Identification of a quality-control mechanism for mRNA 5’-end capping. Nature 467, 608–611

59. Varshney, D., Lombardi, O., Schweikert, G., Dunn, S., Suska, O., and Cowling, V. H. (2018) mRNA Cap Methyltransferase, RNMT-RAM, Promotes RNA Pol II-Dependent Transcription. Cell Rep 23, 1530–1542

60. Amberg, D. C., Burke, D. J., Burke, D., Strathern, J. N., and Laboratory, C. S. H. (2005) Methods in Yeast Genetics: A Cold Spring Harbor Laboratory Course Manual, Cold Spring Harbor Laboratory Press

61. Longtine, M. S., McKenzie, A., 3rd, Demarini, D. J., Shah, N. G., Wach, A., Brachat, A., Philippsen, P., and Pringle, J. R. (1998) Additional modules for versatile and economical PCR-based gene deletion and modification in Saccharomyces cerevisiae. Yeast 14, 953–961

62. Bradford, M. M. (1976) A rapid and sensitive method for the quantitation of microgram quantities of protein utilizing the principle of protein-dye binding. Analytical biochemistry 72, 248–254

63. Ramirez, F., Dundar, F., Diehl, S., Gruning, B. A., and Manke, T. (2014) deepTools: a flexible platform for exploring deep-sequencing data. Nucleic Acids Res 42, W187–191

